# Investigating the effects of radiation, T cell depletion, and bone marrow transplantation on murine gut microbiota

**DOI:** 10.1101/2023.07.10.547212

**Authors:** Jakub Kreisinger, James Dooley, Kailash Singh, Dagmar Čížková, Lucie Schmiedová, Barbora Bendová, Adrian Liston, Alena Moudra

**Affiliations:** Department of Zoology, Faculty of Science, Charles University, Prague, Czech Republic; Immunology Programme, The Babraham Institute, Cambridge CB22 3AT, UK; Department of Medical Cell Biology, Uppsala University, 751 05 Uppsala, Sweden; Institute of Vertebrate Biology, Czech Academy of Sciences, Brno, Czech Republic

## Abstract

Microbiome research has gained much attention in recent years as the importance of gut microbiota in regulating host health becomes increasingly evident. However, the impact of radiation on the microbiota in the murine bone marrow transplantation model is still poorly understood. In this paper, we present the major conclusions of our investigation into the effects of radiation and subsequent bone marrow transplantation with or without T cell depletion of the donor bone-marrow on the microbiota of the ileum and cecum. Our findings show that radiation has different effects on the microbiota of the two intestinal regions, with the cecum showing increased interindividual variation, suggesting an impaired ability of the host to regulate microbial symbionts, consistent with the Anna Karenina principle. Additionally, we observed changes in the ileum composition, including an increase in bacterial taxa that are important modulators of host health, such as *Akkermansia* and *Faecalibaculum*.

In contrast, radiation in the cecum was associated with an increased abundance of several common commensal taxa in the gut, including *Lachnospiraceae* and *Bacteroides*. Finally, we found that high doses of radiation had more substantial effects on the caecal microbiota of the T-cell-depleted group than that of the non-T-cell-depleted group. Overall, our results contribute to a better understanding of the complex relationship between radiation and the gut microbiota in the context of bone marrow transplantation and highlight the importance of considering different intestinal regions when studying microbiome responses to environmental stressors.

## Introduction

The gut microbiota has a massive impact on host health and affects outcomes of various medical treatments through the production of a wide range of bioactive molecules and/or interactions with host tissues and the immune system. Despite extensive research on the changes in the gut microbiota following exposure to ionizing radiation, the microbiota’s specific role in murine bone marrow transplantation (BMT) remains poorly understood.

When exposed to ionizing radiation, certain tissues, including the gut epithelia, which have a high rate of cell proliferation, are particularly vulnerable to damage^1^. This damage often triggers changes in the symbiotic microbiota toward a pathological phenotype, which may amplify the deleterious effects of radiation. On the other hand, the presence of specific microbiota species capable of mitigating radiation-induced tissue damage by producing anti-inflammatory molecules predicts better survival and recovery after irradiation^2^. The effect of radiation has been studied primarily on bacterial communities inhabiting the lower intestine (i.e., caecum and colon). However, little is known about the effects of radiation on the gut microbiota outside the lower gut, which has a different taxonomic composition^3^ and mediates specific functions for the host^4,5^.

Intestinal microbiota plays a crucial role in developing graft-versus-host disease (GVHD) and hematopoietic stem cell transplantation (HCT) outcomes. Studies using allogeneic bone marrow transplantation (BMT) mouse models have characterized the exaggerated inflammatory mechanisms that lead to acute GVHD in target organs^6,7^. These studies have demonstrated that alterations in the composition and diversity of the gut microbiota can disrupt immune homeostasis and contribute to the pathogenesis of GVHD. One common way to prevent GVHD is T-cell depletion of the donor bone marrow cells^8^. Most early trials documented that T-cell depletion could substantially limit acute and chronic GVHD^9^. While there is no direct evidence linking T-cell depletion to changes in the microbiota, it is known that GVHD, which can be prevented by T-cell depletion, can cause severe inflammation and damage to the gut^10^. Overall, the impact of T-cell depletion during bone marrow transplantation on the microbiota is an area that requires further research.

The gut microbiota has been shown to modulate the human immune system, and its impact on HCT outcomes and transplant rejection risk has been increasingly recognized^11,12^. Research in human cohorts has revealed that the composition of the gut microbiota prior to transplantation can influence the likelihood of GVHD development and the overall success of the transplant. Specifically, dysbiosis, characterized by an imbalance in the relative abundance of microbial species, has been associated with increased GVHD incidence and reduced survival rates following transplantation. Recent studies have also highlighted the emerging role of gut microbiota and its metabolites in profoundly impacting allogeneic hematopoietic stem cell transplantation^13^. These studies have shown that the gut microbiota can influence the efficacy of the transplantation process, including engraftment and immune reconstitution^14^. Moreover, specific microbial metabolites, such as short-chain fatty acids, have been implicated in regulating immune responses and promoting the tolerance of donor cells.

To address the above knowledge gaps, we designed an experiment to evaluate the effects of radiation and T-cell depletion separately. By exposing subjects to different doses of radiation, we also investigated the mutual interactions between radiation and T-cell depletion on the resulting composition of the gut microbiota. Last but not least, in addition to the microbiota of the lower intestine represented by caecum samples, we also analyzed the microbiota communities in the ileum. In this way, we were able to gain a unique insight into the effects of BMT (and ionizing radiation in general) on the microbiota of the small intestine. We present the major conclusions of our investigation into the effects of radiation and subsequent bone marrow transplantation with or without T cell depletion on the microbiota of the ileum and caecum using the congenic bone marrow model C57Bl6/J mouse model.

## Main findings

- Combined effect of radiation and antibiotics treatment has different effects on the microbiota of the ileum and the cecum.
- In the cecum, interindividual variation increases due to the combined effect of antibiotics treatment, radiation and BMT, indicating the impaired ability of the host to regulate microbial symbionts (also known as the Anna Karenina principle^15^)
- Changes in ileum composition include bacterial taxa that are important modulators of host health: *Akkermansia* (anti-inflammatory effect^16^) *Faecalibaculum* (antitumor effect^17^)
- In the cecum, radiation is associated with an increased abundance of common commensal taxa in the gut, e.g., *Lachnospiraceae* and *Bacteroides*.
- High doses of radiation have more substantial effects on the caecal microbiota of the T cell-depleted group than that of the non-T cell-depleted group.

## Methods

### Experimental animals

C57BL/6.SJL-*Ptprc*^*a*^/BoyJ (CD45.1) mice were bred and maintained in the Babraham Institute Biological Support Unit (BSU).

No primary pathogens or additional agents listed in the FELASA recommendations^18^ were detected during health monitoring surveys of the stock holding rooms. The ambient temperature was ∼19–21°C, and the relative humidity was 52%. Lighting was provided on a 12 h light: 12 h dark cycle, including 15□min ‘dawn’ and ‘dusk’ periods of subdued lighting. After weaning, mice were transferred to individually ventilated cages with 1–5 mice per cage. Mice were fed CRM (P) VP diet (Special Diet Services) ad libitum and received seeds (e.g., sunflower, millet) at the time of cage cleaning as part of their environmental enrichment. The Babraham Institute Animal Welfare and Ethical Review Body approved all the mouse experimentation. Animal husbandry and experimentation complied with existing European Union and United Kingdom Home Office legislation and local standards (PPL: PP3981824). Sample sizes for mouse experiments were chosen in conjunction with the ethics committees to allow for robust sensitivity without excessive use. Males and females were used for experiments. Age- and sex-matched pairs of animals were used in the experimental groups. Where possible, littermates were equally divided into the experimental groups.

### Bone marrow chimaeras

For the bone marrow (BM) reconstitution experiment, CD45.1 mice were exposed to varying doses of radiation (refer to Table S1 for detailed doses) and subsequently intravenously reconstituted with 2–5 × 10^6^ freshly-collected total BM cells from C57Bl6 donor mice. The BM cells were obtained by crushing the tibia and fibula using a mortar and pestle, followed by filtration through a 50-um filter and red blood cell (RBC) lysis. The total cell count was determined using a hemocytometer. The resulting BM mixture was subjected to T cell depletion using CD4 and CD8 T cell MicroBeads (Miltenyi, cat. No. 130-117-043 and 130-117-044) or injected without T cell depletion.

To prevent infections, the experimental animals undergoing bone marrow transplantation (BMT) were treated with the fluoroquinolone-type antibiotic enrofloxacin (Baytril, Bayer) (50 mg/ml) in their water bottle for the initial three weeks after irradiation, following the standard protocol. The mice were allowed 10 weeks after BMT for BM reconstitution, providing a 7-week window for microbiome reconstitution. The well-being of the experimental animals was monitored daily, and mice that exhibited a weight loss of 20% or showed moderate signs of stress (such as intermittent hunching, piloerection, reduced activity, or poor grooming) were euthanized on ethical grounds. Animals affected by the treatment procedure were excluded from the final dataset.

Upon completion of the experiment, the animals were euthanized, and samples from the duodenum, jejunum, ileum, cecum, and colon were collected for high-throughput 16S rRNA sequencing of the intestinal microbiota.

### Sample preparation

Experimental animals were euthanized, and the colon, caecum, and ileum were sampled, including both intestinal walls and contents for 16S barcoding. The samples were stored in 2 ml ethanol (VWR, 20821.330P). Clean and fire-sterilized scissors and forceps were used for each intestinal section to minimise bacterial DNA contamination. A negative control cryotube was prepared for each round of dissections to sample the possible air-borne contamination. The ileum, caecum, and colon were sampled separately, with the contents and walls placed in separate cryotubes and frozen at -80°C. An additional faecal sample was taken from the last portion of the colon.

### Sample processing

Metagenomic DNA was extracted using the PowerSoil kits (QIAGEN). Gut microbiome sequencing libraries were prepared using the two-step PCR protocol at the Studenec Research Facility of the Institute of Vertebrate Biology, CAS, as described in detail in Čížková et al. (2021)^19^. Briefly, standard metabarcoding primers for the V3-V4 hypervariable region of bacterial 16S rRNA^20^ were used in the first PCR step to amplify the specific rRNA loci. Dual indexes were introduced in the second PCR, during which Illumina-compatible Nextera-like sequencing adaptors were reconstituted. Each metabarcoding PCR was performed in a technical duplicate to account for the stochasticity of PCR and sequencing. PCR products were pooled according to their concentration, and pools were sequenced using Illumina Miseq (v3 kit, 300 bp paired-end reads) at CEITEC, Brno, Czech Republic.

### Bioinformatics

Skewer^21^ was used to demultiplex samples and detect and trim gene-specific primers. In the next step, low-quality reads (expected error rate per paired-end read > 2) were eliminated, quality-filtered reads were denoised, and abundance matrices were generated using the read counts for each 16S rRNA amplicon sequencing variant (hereafter ASVs) in each sample. All these steps were performed using the R package dada2^19^. Subsequently, uchime^22^ was used in conjunction with the gold.fna reference database (available at: https://drive5.com/uchime/gold.fa) to detect and eliminate chimeric ASVs. The taxonomy for the non-chimeric ASVs was assigned with 80% posterior confidence by the RDP classifier^23^ using the Silva database v.138^24^ as a reference. Procrustean analysis revealed high consistency between technical duplicates (r = 0.97, p = 0.0001). Consequently, we merged the microbiota profiles of the technical duplicates, eliminating all ASVs that were not detected in both duplicates. Finally, we excluded all samples with a number of high-quality reads < 1000. The final dataset included data from 82 mice whose gut microbiota was sequenced. We were unable to sequence two ileum samples due to insufficient concentration of template DNA, so the final data set included a total of 182 samples (82 from the caecum and 80 from the ileum). The total number of sequences for the entire data set was 1 599 231, with an average sequencing coverage per sample of 9872 (range = 1532 - 19507). A total of 1016 bacterial ASVs were detected. Sample metadata for this study is shown in **Table S1**.

The final dataset was used for metagenome predictions using the picrust2 pipeline^25^ with default parameters, where predicted metagenomes were categorized into functional pathways^26^. Using the BugBase pipeline, we also estimated the frequency of selected phenotypic traits for each sample, including, for example, stress and oxygen tolerance, pathogenicity potential, biofilm formation ability, or Gram positivity/negativity^27^.

### Statistics

The microbiota of the caecum was clearly different from the microbial profiles of the ileum (**Figure S1**). Therefore, we analyzed these two sample types separately. For the entire data set, we first tested whether the microbiota differed between experimental groups (non-irradiated controls vs irradiated depleted and irradiated non-T cell-depleted mice) using mixed effects models. The effect of sex and experimental replicate were included as covariates, and cage identities as random effects. In the subsequent analysis step, we removed the control individuals (who were not irradiated and did not receive antibiotics treatment, n = 12) from the data to examine the effect of radiation dose. Because of the dichotomous distribution of radiation doses, in which mice were exposed to relatively low doses between 4.5 and 5.5 Gy (n = 24) or high radiation doses between 10 and 13 Gy (n = 46), we considered these two categories as a factorial predictor. Apart from the effect of radiation dose and the set mentioned above of covariates, we also tested whether the effect of radiation dose differed between the depleted and non-T-cell-depleted groups by including an interaction term in these statistical models.

Statistical analyses considered three levels of microbiota variation. First, we focused on alpha diversity variation, assessed based on the Shannon diversity index and ASV richness (i.e., the total number of ASVs) calculated for each sample. These alpha diversity indices were considered response variables in mixed models with Gaussian error distribution fitted with the R package lme4. To achieve a normal distribution of the residuals, log_10_-transformed values of ASV richness were used.

Next, variation in microbiota composition was analyzed using mixed models from the R package MDMR^28^, with compositional dissimilarity between samples as the response. Two dissimilarity indices were considered: the Jacccard index, which considers only variation in the presence/absence of ASV, and the Bray-Curtis index, which also considers variation in relative ASV abundances. We focused on systematic shifts in the community and analyzed differences in interindividual dispersion between experimental groups. For these analyses, we calculated Euclidean distances to centroids of each of the three experimental groups using the betadisper function from the R package vegan. Consequently, individuals with highly divergent microbiota content compared with other individuals within the same experimental group received high scores. Distances to centroids were log_10_-transformed and then used as response variables in mixed models fitted with the R package lme4. Finally, we examined variation in the relative abundance of individual ASVs using mixed models with negative binomial distribution fitted with the R package glmmTMB, using the number of reads for each ASV in each sample as the response variable. To avoid false positives due to multiple testing, we calculated false discovery rates (FDR)^29^ based on the resulting probability values and considered only those results as significant where the FDR was less than 0.05.

To account for differences in sequencing depth between samples, we included log-transformed read counts for each sample at the model offset in the ASV-level analyses. In the case of the alpha-diversity and dissimilarity-based analyses, diversity indices were calculated based on rarefaction datasets, with the rarefaction threshold corresponding to the minimum sequencing depth achieved.

Statistical analyses of metagenome predictions followed a similar logic, with two exceptions. First, alpha diversity analyses were not performed for the predicted metagenomes. Second, we did not consider absence/presence-based Jaccard dissimilarity, as it is less informative given the functional redundancy of microbial communities.

## Results

### Variation in alpha diversity

For the entire data set of microbial profiles of the cecum and ileum, we did not detect any effects of radiation or radiation combined with depletion on microbial alpha diversity. The same was true for all other predictors tested, including the experimental replicate (**Table S2**). After excluding control mice from the data set, significant interactions between depletion treatment and radiation dose indicated a differential suggested divergent effect of radiation intensity on alpha diversity of caecum microbiota in depleted versus non-T cell-depleted mice. However, there were no effects of radiation dose or other covariates on the ileum microbiota (**Table S3**). Separate models for non-T cell-depleted mice showed an increase in alpha diversity of the caecal microbiota in non-T-cell-depleted mice exposed to higher radiation doses (ASV richness: Estimate = 0.0550 ± 0.0246 [± SE], ΔD.f. = 1, χ^2^ = 5.2462, p = 0.0220, Shannon: Estimate = 0.2237 ± 0.0837 [± SE], ΔD.f. = 1, χ^2^ = 7.89, p = 0.0050), whereas there was no such difference in depleted individuals (ASV richness: Estimate = −0.0240 ± 0.0387 [± S.E.], ΔD.f. = 1, χ^2^ = 0.5859, p = 0.4440, Shannon: Estimate = -0.1391 ± 0.1223 [± SE], ΔD.f. = 1, χ^2^ = 1.9432, p = 0.1633).

### Variation in composition

After statistically controlling for variation between experimental replicates and other confounding factors, both irradiated groups’ caecum and ileum microbiota showed significant compositional changes compared with non-irradiated controls without antibiotics (**Table S4, Figure 1**). This was also confirmed by additional MDMR analyses aimed at pairwise comparisons of all treatment groups, which consistently revealed significant differences between controls and depleted mice or controls and non-T-cell-depleted mice (p < 0.01 in all cases). In contrast, differences between depleted and non-T-cell-depleted mice were insignificant (p > 0.1 in all cases). In addition to the effects of radiation, antibiotics and BMT on microbiota composition, both MMDR analyses and PCoA-based sample ordination (**Figure 1, Table S4**) showed highly significant changes between experimental replicates. While the microbiota of the control group was comparable in both replicates, the microbiota of both irradiated mice showed a pronounced divergence greater than the variation between irradiated and non-irradiated mice within the experimental replicates (**Table S4**).

**Figure 1.**
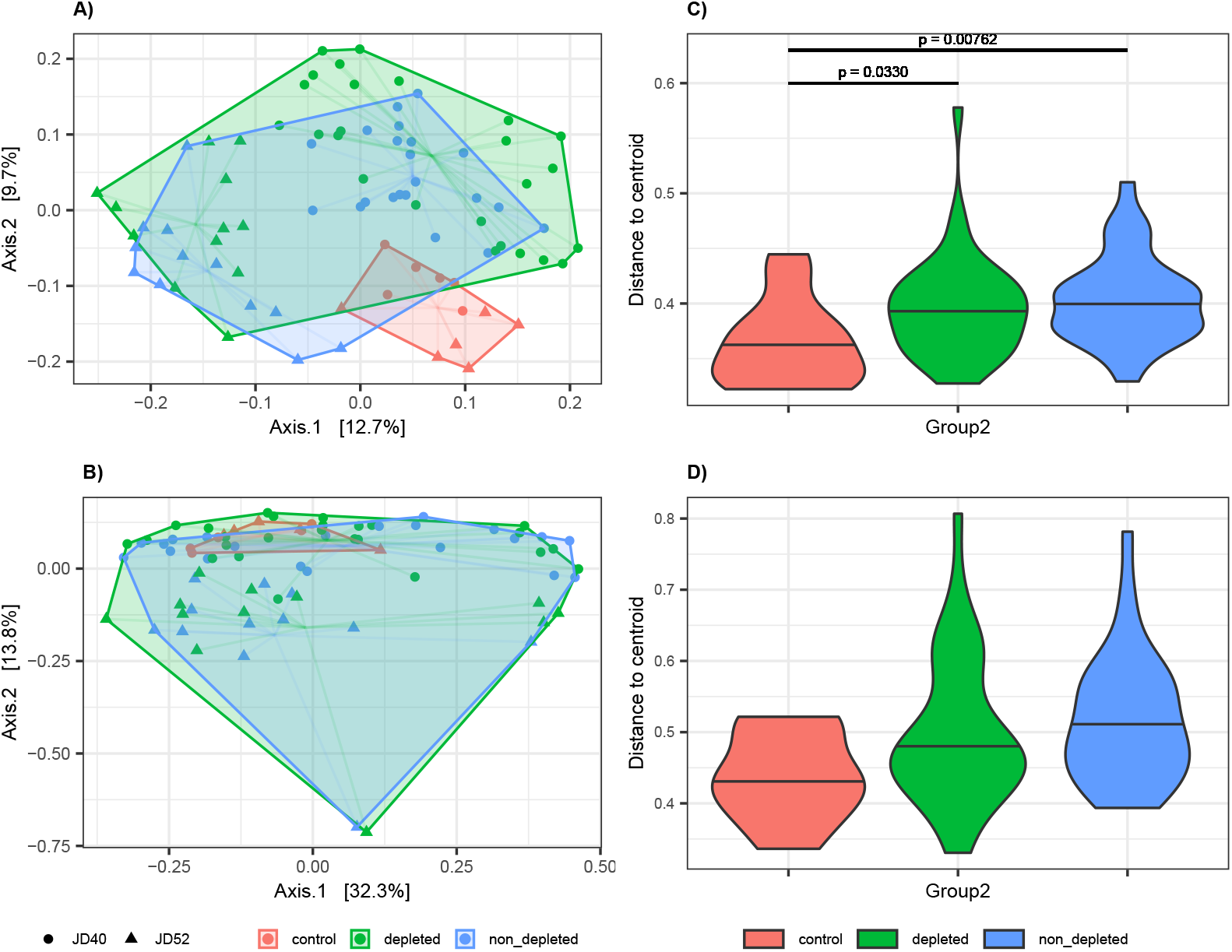
[BETA_all.ja.pdf]: Variation in microbiota composition between non-irradiated controls and depleted or non-T-cell-depleted mice exposed to radiation. Shifts in the composition of the microbiota of A) cecum and B) ileum between treatment groups were analyzed by PCoA ordination of Jaccard dissimilarities. Different colours indicate treatment levels. Harwestman diagrams link samples from the same treatment replicate. Interindividual variation for each treatment group is shown for C) caecal and D) ileum samples by violin plots showing the distribution of Jaccard-based distances to group-specific centroids. Horizontal lines within violin plots correspond to median values. Horizontal lines above the violin plots indicate significant differences between the experimental groups.

Apart from the systematic compositional shifts due to radiation and subsequent course of enrofloxacin antibiotic treatment revealed by MDMR analyses, PCoA (**Figure 1**) also suggested that the irradiated mice exhibited increased interindividual dispersion, as indicated by a larger ordination space occupied. This pattern may result from perturbed mechanisms regulating the symbiotic microbiota. The administration of a broad-spectrum antibiotic enrofloxacin causes a state of dysbiosis in which there is an elevated proportion of *Clostridium coccoides, C. coccoides-Eubacterium rectale, Bacteroidetes*, and *Bifidobacterium spp*, while the presence of segmented filamentous bacteria is reduced.

To statistically test this type of variation, linear mixed models were fitted, where Euclidean distances to PCoA centroids for sample groups determined by the combination of treatment level and experimental repetition were included as a response. For the Jaccard index and caecum microbiota, this approach revealed increased interindividual dispersion of both irradiated groups compared with the control (**Figure 1, Table S5**). In contrast, for the Bray-Curtis dissimilarities, the difference between the controls and T-cell-depleted mice was only marginally significant (p = 0.0799). In the case of ileum microbiota, the differences in interindividual dispersion between experimental groups were only marginally significant when Jaccard dissimilarities were considered and not significant for Bray-Curtis dissimilarities (**Table S5**).

Additional analyses were performed for a data subgroup without non-irradiated controls to determine whether the microbiota changed with radiation dose and whether this change was modulated by depletion treatment. There was only a mild significant effect of radiation dose on the ileum microbiota when Jaccard dissimilarities were used, and there was an effect on the ileum microbiota based on Bray-Curtis dissimilarities. However, a highly significant interaction between depletion treatment and radiation dose in the caecum samples suggested a differential effect of radiation intensity in T-cell-depleted versus non-T-cell-depleted mice (**Table S6**). In a separate analysis for the subset of caecum samples from the T-cell-depleted mice, the effect of radiation dose was highly significant (MDMR: D.f. = 1, MDMR statistic = 20.7289, p < 0.0001 for Bray-Curtis and D.f. = 1, MDMR statistic = 5.7491, p < 0.0001 for Jaccard dissimilarities) and that the same was true for non-T-cell-depleted groups (MDMR: D.f. = 1, MDMR statistic = 4.8182, p < 0.0001 for Bray-Curtis and D.f. = 1, MDMR statistic = 1.881629, p = 0.0172 for Jaccard dissimilarities), although the effect size was much smaller. This was consistent with the PCoA sample ordination (**Figure 2**), in which the depleted mice exposed to high and low doses of radiation formed almost no overlapping clusters. In contrast, considerable overlap was observed in the non-T-cell-depleted group exposed to different doses of radiation.

**Figure 2.**
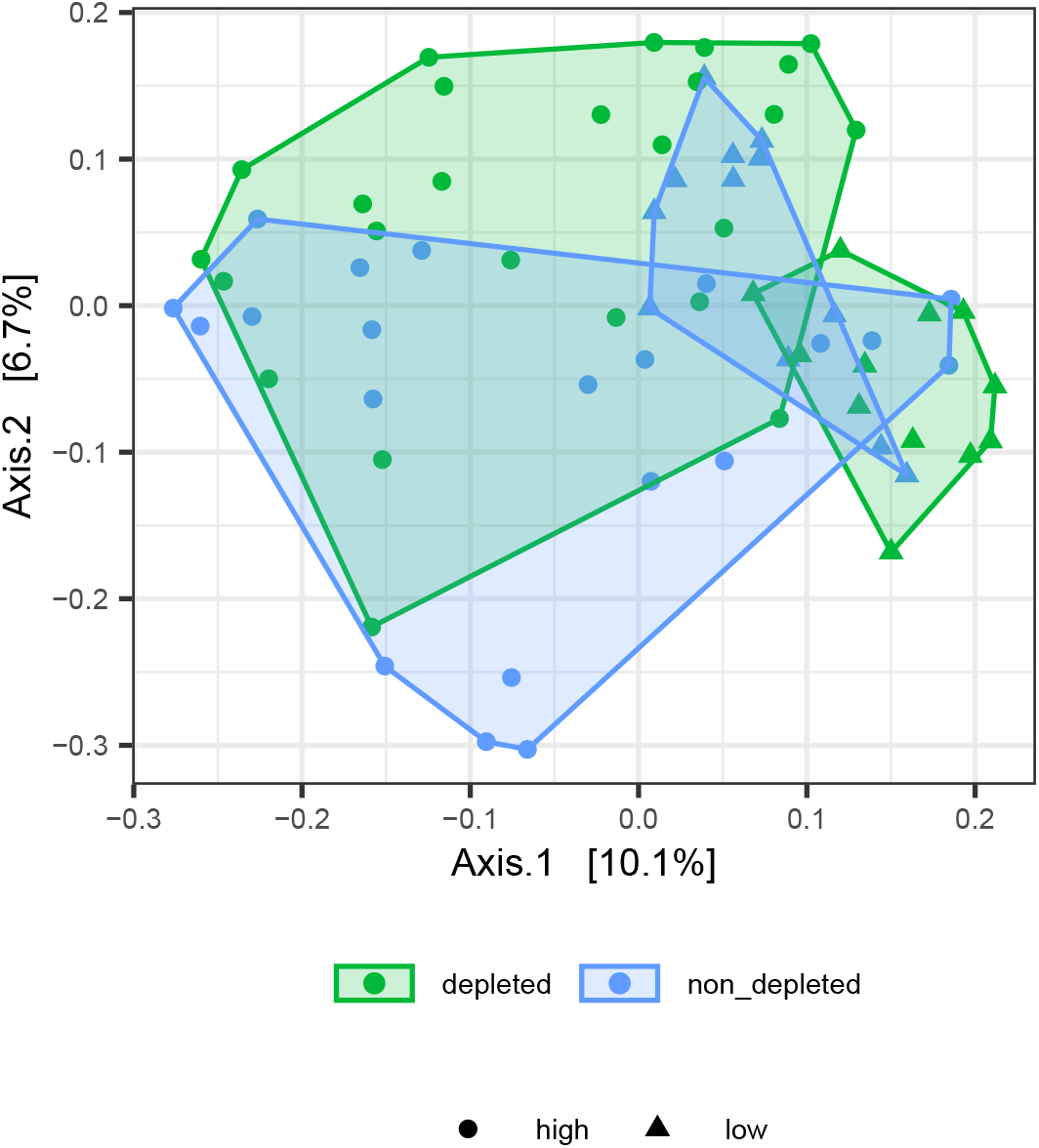
[Jac_dose.pdf]: PCoA ordination of Jaccard dissimilarities depicts variation in caecum microbiota composition between depleted and non-T-cell-depleted mice exposed to either high or low doses of radiation.

### Variation in ASVs relative abundances

Differential abundance analyses comparing the caecal microbiota of the non-irradiated controls with that of the two irradiated groups identified five ASVs from the genus *Bacteroides* and the family *Lachnospiraceae* whose relative abundance was increased in the irradiated mice. Complementary analyses for the ileum microbiota revealed a decreased abundance of *Faecalibaculum* and *Muribaculum* and an increased abundance of *Akkermansia* in the mice exposed to radiation and subsequent antibiotics treatment (**Figure 3**).

**Figure 3.**
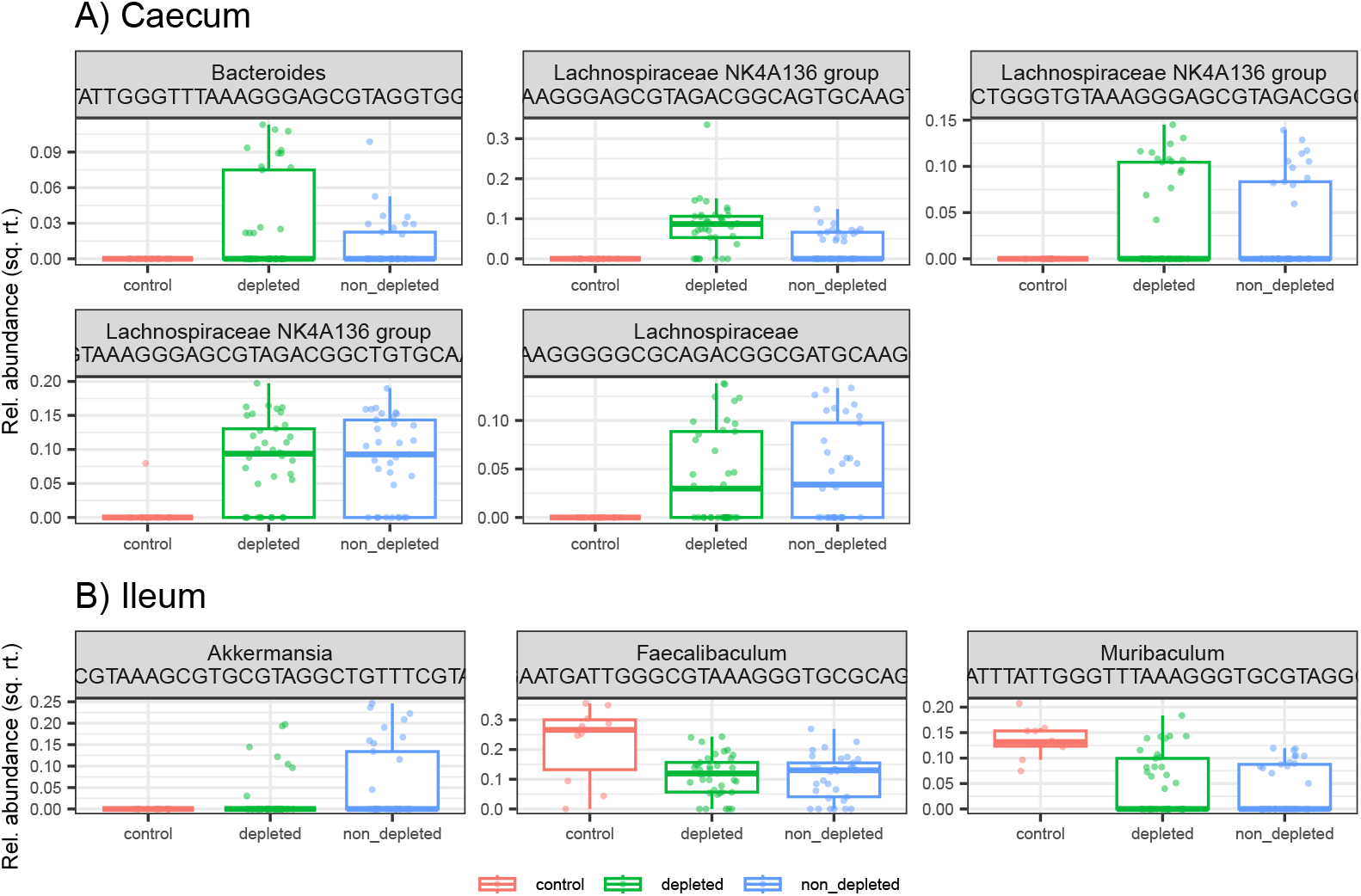
**[DFA_all.pdf]**: Effect of radiation on ASVs relative abundances. Box plots showing a variation of ASVs whose relative abundance in A) caecum and B) ileum microbiota differs among three treatment groups represented by non-irradiated controls, irradiated depleted mice, and irradiated non-T-cell-depleted mice.

**Figure 4.**
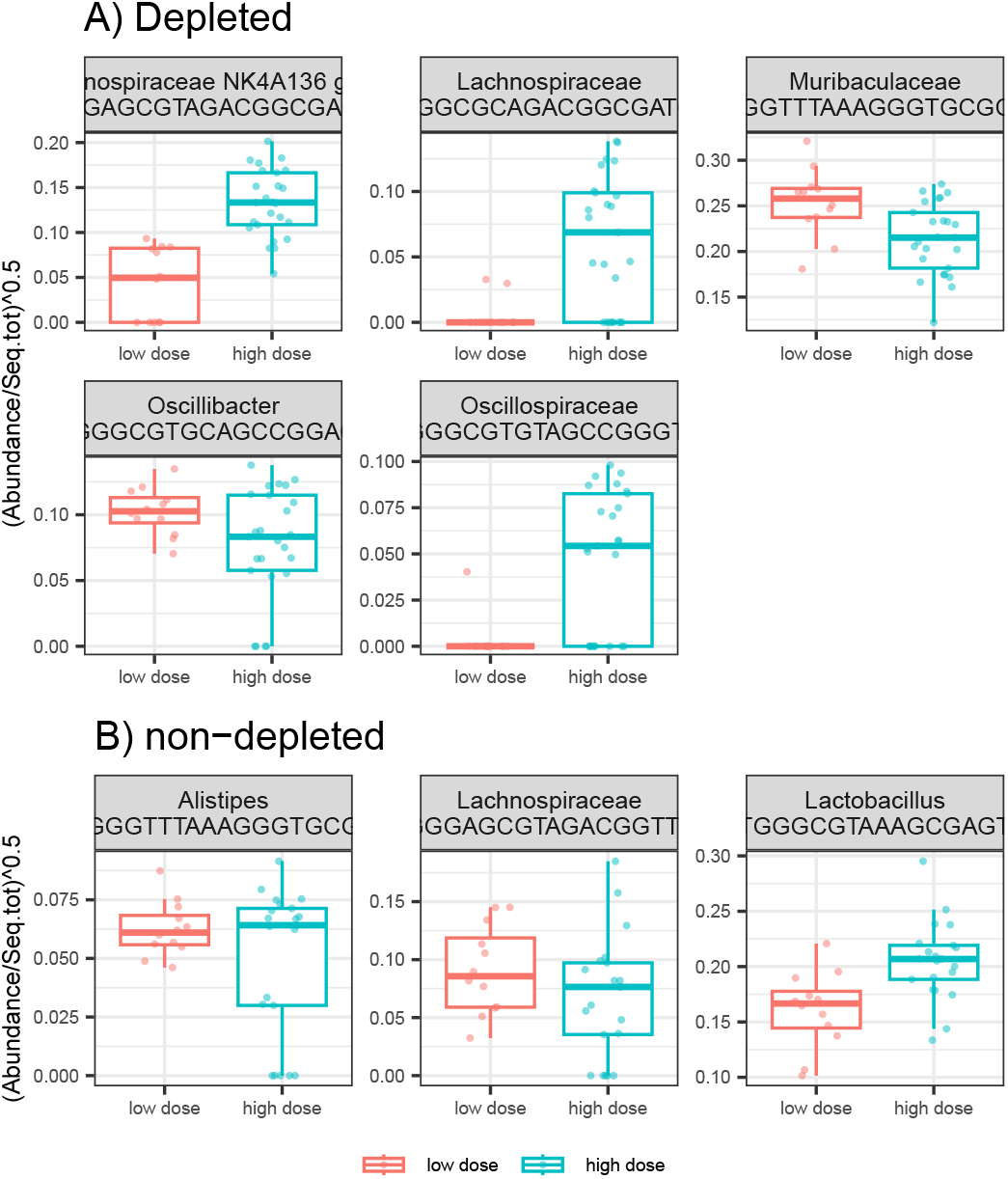
[DFA_depl_nondepl.pdf]: Effect of radiation dose and depletion treatment on ASVs relative abundances. Boxplots showing the variation of ASVs whose relative abundance in the cecum of A) the depleted group and B) the non-T-cell-depleted group differed between individuals exposed to a low and a high radiation dose.

Differential abundance analyses for the caecal microbiota, testing the effect of radiation dose, revealed different ASVs variation patterns for the T-cell-depleted and non-T-cell-depleted mice. For the subgroup of T-cell-depleted mice, high radiation doses resulted in an increased abundance of ASVs from *Lachnospiraceae* and *Oscillospiraceae*, whereas ASVs from the genus *Oscillospira* and the family *Muribaculaceae* showed the opposite direction of change. On the other hand, non-T-cell-depleted mice exposed to high radiation and subsequent antibiotics treatment showed an increased abundance of ASVs from the genus *Lactobacillus* and decreased abundance of ASVs from the genus *Alistipes* and *Lachnospiraceae*.

### Predicted microbiota functions

According to MDMR analyses, the content of caecal metagenomes differed between non-irradiated controls and depleted (Tukey posthoc tests: p < 0.0001) or non-T-cell-depleted (Tukey posthoc tests: p = 0.0004) mice, both of which were exposed to radiation. However, experimental groups had no significant differences in predicted metagenomes of the ileum microbiota (**Table S4, Figure S4**). This was consistent with differential abundance analyses identifying 14 metabolic pathways, whose predicted abundances differed between controls and irradiated groups (**Figure S6**). Interindividual variation did not vary significantly between experimental groups (**Table S5**), although PCoA indicated increased interindividual variation in mice exposed to radiation (**Figure S4**). MDMR for a data set without non-irradiated controls showed that radiation dose significantly altered the caecum microbiome but not the ileum (**Figure S5, Table S5**). In addition, consistent with the same analyses for microbiota composition, the interaction between radiation dose and depletion treatment was significant for caecum microbiota. Differential abundance analyses identified five predicted pathways for depleted and 34 pathways for non-T-cell-depleted groups, whose expression varied with radiation dose. However, estimates characterizing abundance changes between mice exposed to high vs low radiation intensity calculated separately for depleted and non-T-cell-depleted groups were tightly correlated (r = 0.6693, p <0.00001).

BugBase predictions revealed no differences between experimental groups in the proportion of aerobic, facultative aerobic, Gram-negative, stress-tolerant, or potentially pathogenic bacteria and bacteria that can form biofilms (**Figure S8**, LMM: p > 0.05 in all case).

## Discussion

Our study aimed to shed light on the complex relationship between radiation, BM transplantation, and gut microbiota. We specifically focused on the ileum and cecum as these regions have distinct functions and harbour different microbial communities^4,5^. Our findings suggest that radiation and subsequent antibiotic treatment have different effects on the microbiota of these two intestinal regions. In particular, we found that the irradiated cecum exhibited increased interindividual variation and impaired ability of the host to regulate microbial symbionts, consistent with the Anna Karenina principle. These observations suggest that radiation and subsequent antibiotic treatment may significantly impact the stability and functionality of the caecal microbiota.

Our study revealed noteworthy changes in the microbiome composition of the ileum, the last section of the small intestine, following radiation exposure and subsequent antibiotics treatment. Specifically, we found an increase in the abundance of certain bacterial taxa, such as *Akkermansia*, which are crucial in maintaining host health. In contrast, we observed a decrease in the *Faecalibaculum* and *Muribaculum*.

Of particular interest, *Akkermansia* has been shown to have numerous benefits, including enhancing glucose metabolism, promoting intestinal barrier function, and exerting anti-inflammatory effects.^16^ Meanwhile, *Faecalibaculum* has been associated with antitumor properties and the production of short-chain fatty acids that facilitate the production of IgA by plasma cells.^17^ Interestingly, the levels of *Akkermansia* and *Faecalibaculum* in the gut microbiota have been found to positively correlate with the hypoglycemic effects of *Astragalus membranaceus* polysaccharides and the alleviation of food allergy symptoms in mice.^30,31^ Our findings support the crucial role of *Akkermansia* and *Faecalibaculum* in gut microbiota-mediated host health responses and suggest that radiation exposure may impact these important microbial populations.

Comparatively, in humans, after allogeneic BMT, there is a decline in bacterial alpha diversity within the first 3 weeks in the gut, oral, and nasal microbiota.^32–36^ Patients with acute GVHD exhibit the most significant loss of diversity.^34,36–38^ It is noteworthy that both adult and pediatric allo-BMT patients experience reduced diversity and an altered gut bacterial composition even before conditioning and transplantation, compared with healthy individuals.^33,36^ This reduction in diversity may be due to antibiotic treatment, which reduces specific groups of commensal mucosal colonizers. Obligate anaerobes, such as *Clostridiales, Negativicutes, Bacteroidetes*, and *Fusobacteria*, are reported to be especially vulnerable to the adverse effects of Piperacillin-tazobactam and meropenem. In contrast, fluoroquinolones, intravenous vancomycin, and trimethoprim□sulfamethoxazole appear to have a lesser impact on these bacterial groups. ^39,40^

However, we also found that radiation and antibiotics treatment in the caecum was associated with an increased abundance of several common commensal taxa in the gut, including *Lachnospiraceae* and *Bacteroides*. This observation suggests that radiation and subsequent antibiotics treatment may impact the composition of the caecal microbiota differently, depending on the microbial taxa involved.

Finally, we found that high doses of radiation had more substantial effects on the caecal microbiota of the depleted group than that of the non-T-cell-depleted group. This observation highlights the importance of considering the role of immune cells in modulating the gut microbiota in response to environmental stressors.

In conclusion, our study provides valuable insights into the effects of radiation and bone marrow transplantation on the gut microbiota in the murine BM transplantation model. Our findings suggest that radiation and the subsequent antibiotics treatment have different effects on the ileum and caecum microbiota and that immune cells’ role in modulating the gut microbiota should be considered when studying responses to environmental stressors. Further research is needed to fully understand the complex interactions between radiation, gut microbiota, and host health.

## Conclusion

This study investigated the relationship between radiation, bone marrow transplantation (BMT), and gut microbiota in the ileum and caecum. The irradiated caecum showed increased interindividual variation and impaired microbial regulation. Radiation and antibiotics treatment had varying effects on the ileum, with an increase in beneficial bacteria like *Akkermansia* and a decrease in *Faecalibaculum. Akkermansia* and *Faecalibaculum* play important roles in host health. Human BMT patients also experience bacterial diversity and composition changes, particularly in graft-vs-host disease (GVHD) cases. In the caecum, radiation and antibiotics treatment increased the abundance of specific commensal taxa. The impact of radiation on the caecal microbiota was more pronounced in the T-cell-depleted group. Immune cells influence the gut microbiota’s response to environmental stressors. This study provides insights into radiation, BMT, and gut microbiota, highlighting the need for further research to understand their complex interactions.

## Supplementary Tables

**Table S1**: Sample metadata and accession numbers for sequencing data (will be updated upon manuscript acceptance

**Table S2**: Variation in alpha diversity between all experimental groups, i.e., the irrational depleted and non-T-cell-depleted mice and the non-irradiated control mice considered as reference (i.e., intercept). The table shows the estimated parameters of the mixed-effects models, the corresponding standard errors, the test statistic (t), and the resulting significance values (p).

**Table S3**: Variation in alpha diversity due to the effect of radiation dose and other covariates (sex, depletion treatment, and experimental replicate). The table shows the estimated parameters of the mixed-effects models, the corresponding standard errors, the test statistics (t), and the resulting significance values (p).

**Table S4**: Variation in microbiota composition. The effect of treatment level and other covariates was analyzed using mixed MDMR models. The microbiota profiles of the ileum and cecum were analyzed separately using Bray-Curtis and Jaccard dissimilarities. The same models were used to analyze variation in predicted metagenome content, for which Bray-Curtis dissimilarities were used. The table shows the values of the test statistics, the degrees of freedom, and the resulting probability values.

**Table S5**: Variation in the interindividual dispersion of microbiota composition between the treatment groups, i.e., the irradiated depleted and non-T-cell-depleted mice and the non-irradiated control mice. The Euclidean distance to PCoA centroids represented the response variable of these models. Statistical testing was performed by comparing the models with and without the effect of the treatment group as a predictor using likelihood ratio tests. Probability values were derived based on deviance changes assumed to follow the chi-square distribution. Models included all covariates mentioned in the main text, but their effects were always not significant (p > 0.05). A separate model was created for each gut section’s microbiota profiles and predicted metagenomes. Bray-Curtis and Jaccard dissimilarities were used to analyze the microbiota profiles, while only Bray-Curtis dissimilarities were considered in the case of the predicted metagenomes.

**Table S6:** Effect of radiation dose and depletion treatment on the microbiota composition and predicted metagenomes. Data were analyzed using mixed MDMR models. The microbiota profiles of the ileum and cecum were analyzed separately with Bray-Curtis and Jaccard dissimilarities. The same models were used to analyze variation in predicted metagenome content, for which Bray-Curtis dissimilarities were used. The table shows the values of the test statistics, the degrees of freedom, and the resulting probability values.

## Supplementary Figures

**Figure S1 [Caecum_ileum_overview.pdf]: Gut microbiota variation between caecum and ileum**. Consistently for all experimental groups, the ileum microbiota showed lower alpha diversity and different composition compared to the caecal microbiota, as shown by A) violin plots visualizing differences in ASV richness, B) Principal Coordinate analysis for Bray-Curtis dissimilarities, where caecal and ileal samples from non-overlapping clusters and where identity with the experimental group is indicated by Harwestman plots, and by C) bar plots showing the proportions of the 20 most abundant bacterial genera in each sample.

**Figure S2** [BETA_all.bc.pdf]: Variation in microbiota composition between non-irradiated controls and depleted or non-T-cell-depleted mice exposed to radiation. Shifts in the composition of the microbiota of A) cecum and B) ileum between treatment groups were analyzed by PCoA ordination of Bray-Curtis dissimilarities. Different colours indicate treatment levels. Harwestman diagrams link samples from the same treatment replicate. Interindividual variation for each treatment group is shown for C) caecal and D) ileum samples by violin plots showing the distribution of Bray-Curtis-based distances to group-specific centroids. Horizontal lines within violin plots correspond to median values. Horizontal lines above the violin plots indicate significant differences between the experimental groups.

**Figure S3** [Bray_dose.pdf]: PCoA ordination of Bray-Crutis dissimilarities depicts variation in caecum microbiota composition between depleted and non-T-cell-depleted mice exposed to high or low doses of radiation.

**Figure S4** [BETA_all.bc_METAG.pdf]: Variation in predicted metagenome content of A) caecum and B) ileum between treatment groups assessed based on PCoA for Bray-Curtis dissimilarities.

**Figure S5 [DFA_all_METAG.pdf]** Effect of radiation on predicted relative abundances of metabolic pathways. Boxplots show the variation in metabolic pathways whose relative abundance in the caecal microbiota differs among the three treatment groups (non-irradiated controls, irradiated depleted mice, and irradiated non-T-cell-depleted mice). The same analysis for the ileum microbiota revealed no significant differences.

**Figure S6** [Bray_dose_METAG.pdf]: PCoA ordination of Bray Curtis dissimilarities depicts differences in the composition of predicted metagenomes of caecum between depleted and non-T-cell-depleted mice exposed to either high or low doses of radiation.

**Figure S7** [DFA_depl_nondepl_METAG] Effect of radiation dose and depletion treatment on predicted relative abundances of metabolic pathways. Boxplots showing the variation of pathways whose relative abundance in the cecum of A) the depleted group and B) the non-T-cell-depleted group differed between individuals exposed to a low and a high radiation dose.

**Figure S8** [Pred_phenotype.pdf]: Variation in phenotypic traits between experimental groups. Box plots show the proportions of aerobic, facultative aerobic, Gram-negative, stress-tolerant, or potentially pathogenic bacteria and bacteria capable of forming biofilms in the experimental groups and the microbiota samples from the cecum or ileum.

