## Supplementary figures and images for "Investigating the effects of radiation, T cell depletion, and bone marrow transplantation on murine gut microbiota"

### Figure S1

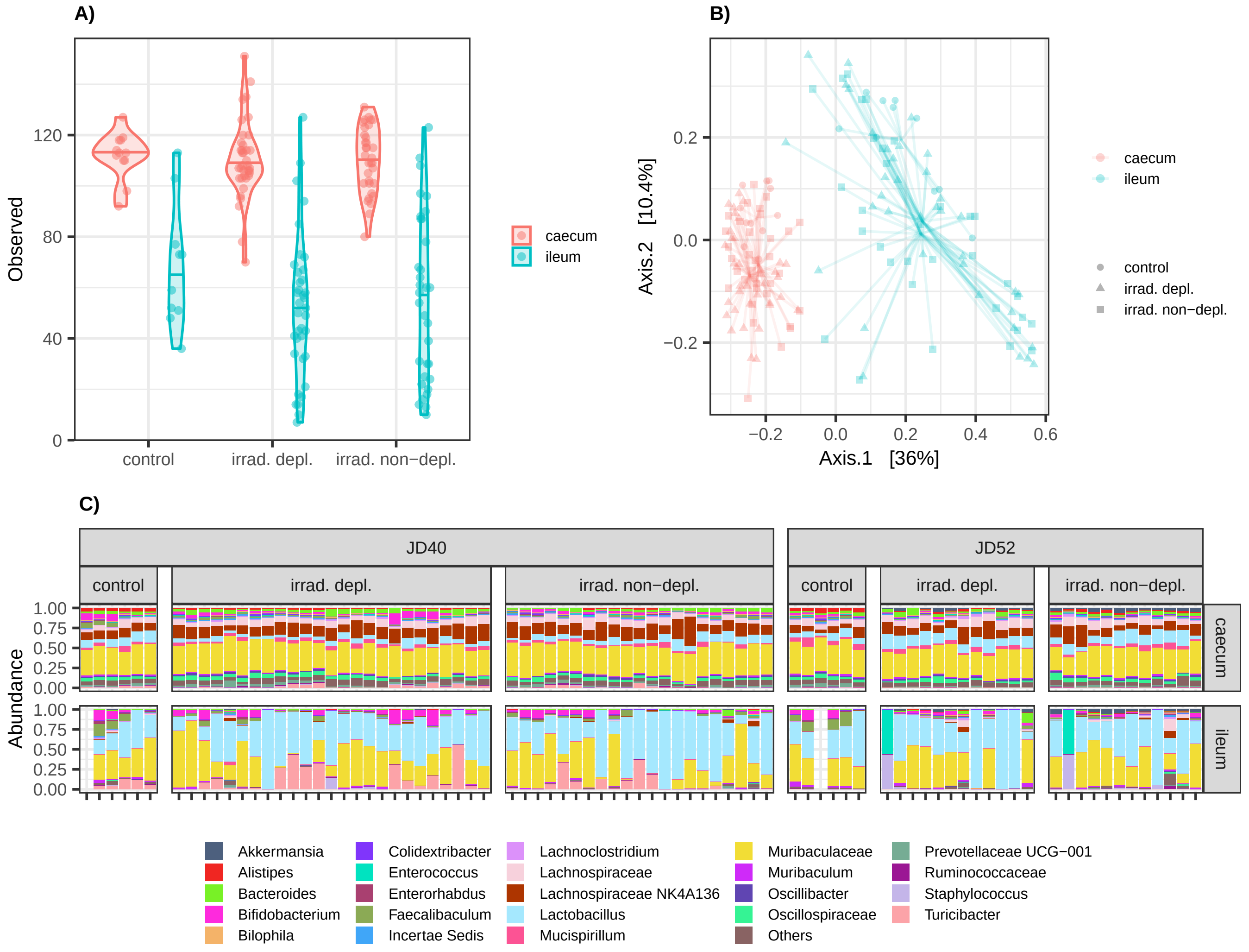

### Figure S2

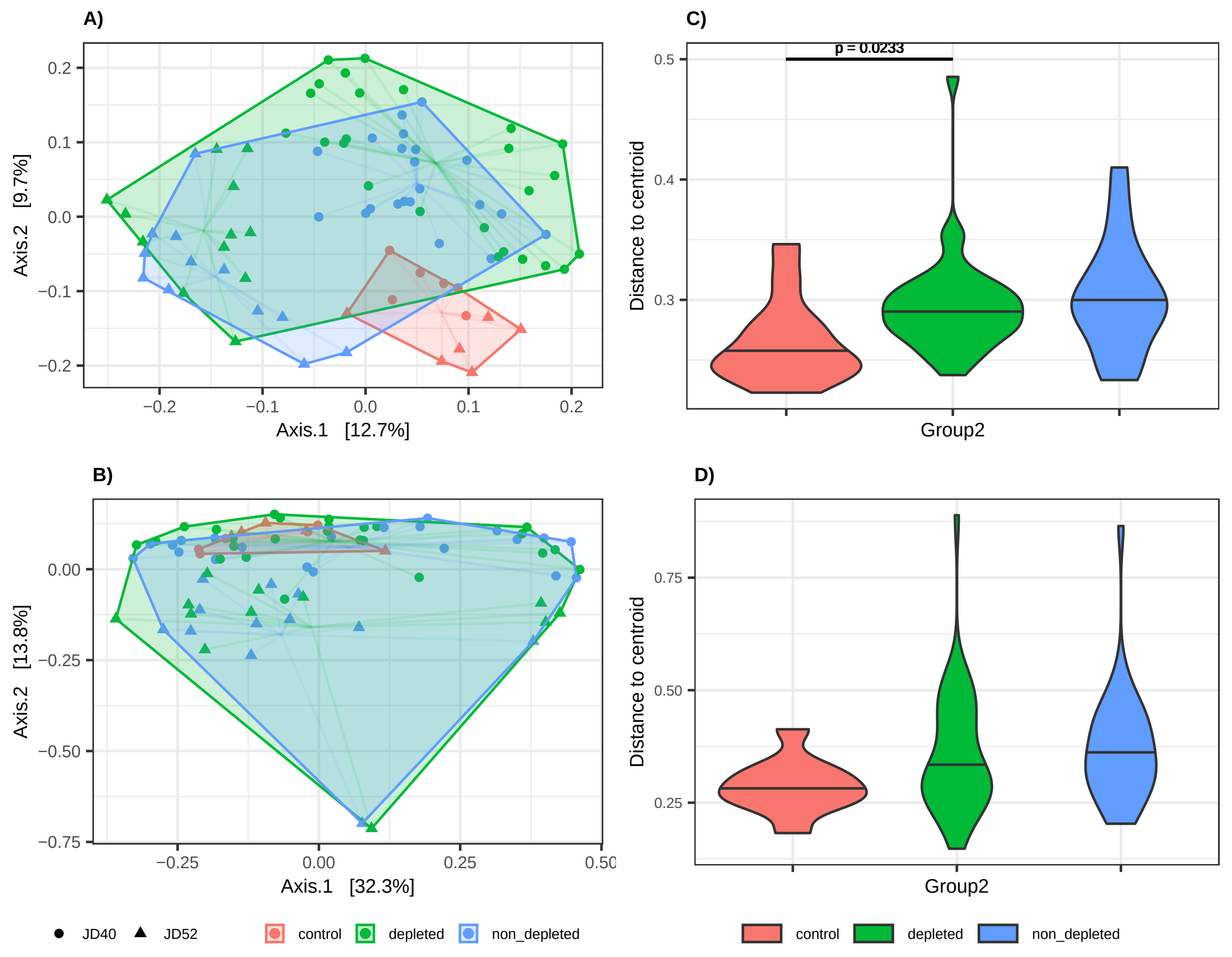

### Figure S3

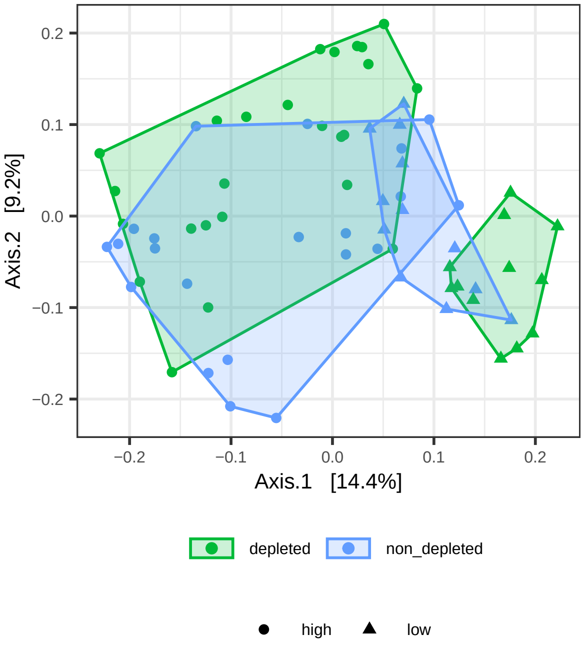

### Figure S4

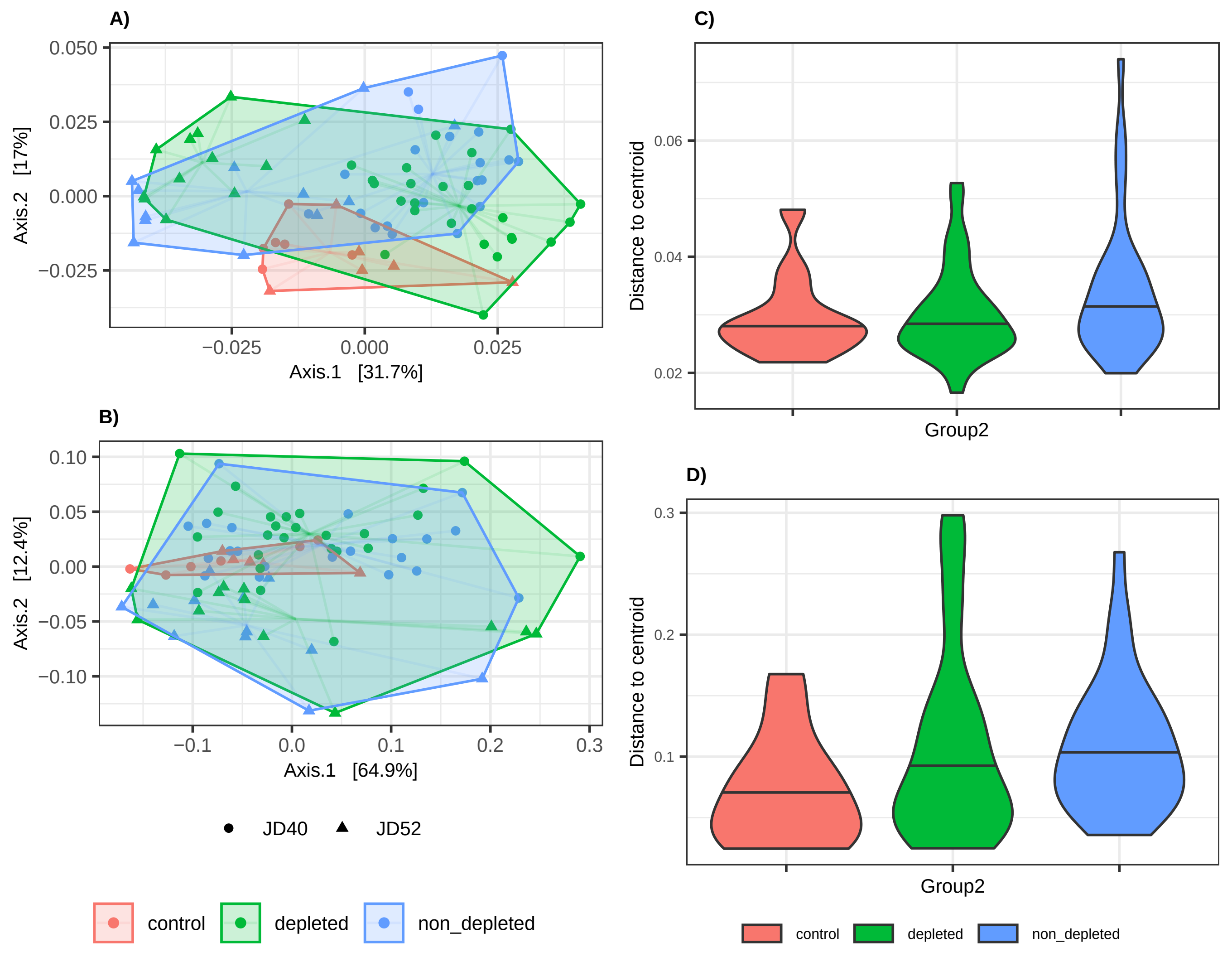

### Figure S5

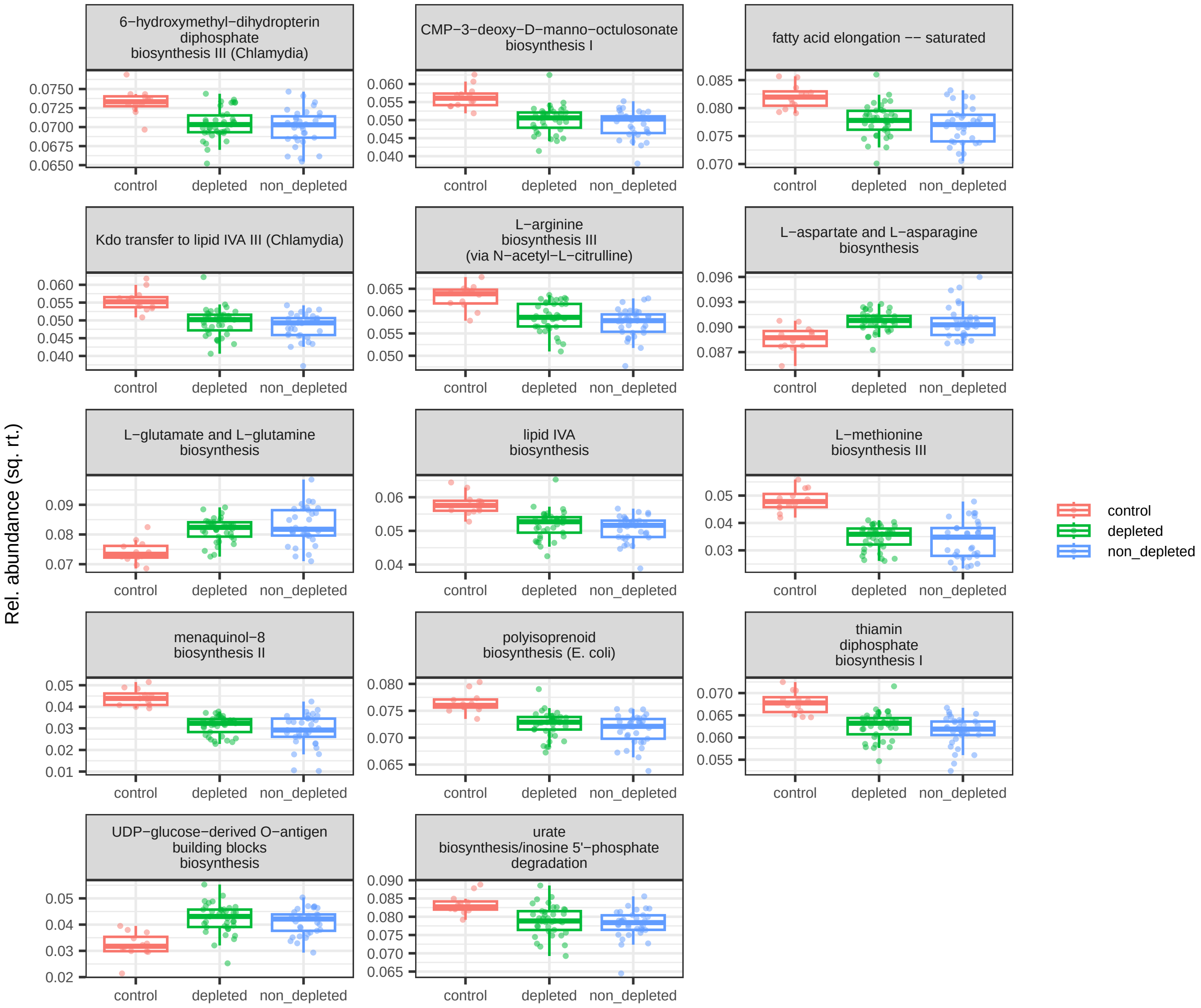

### Figure S6

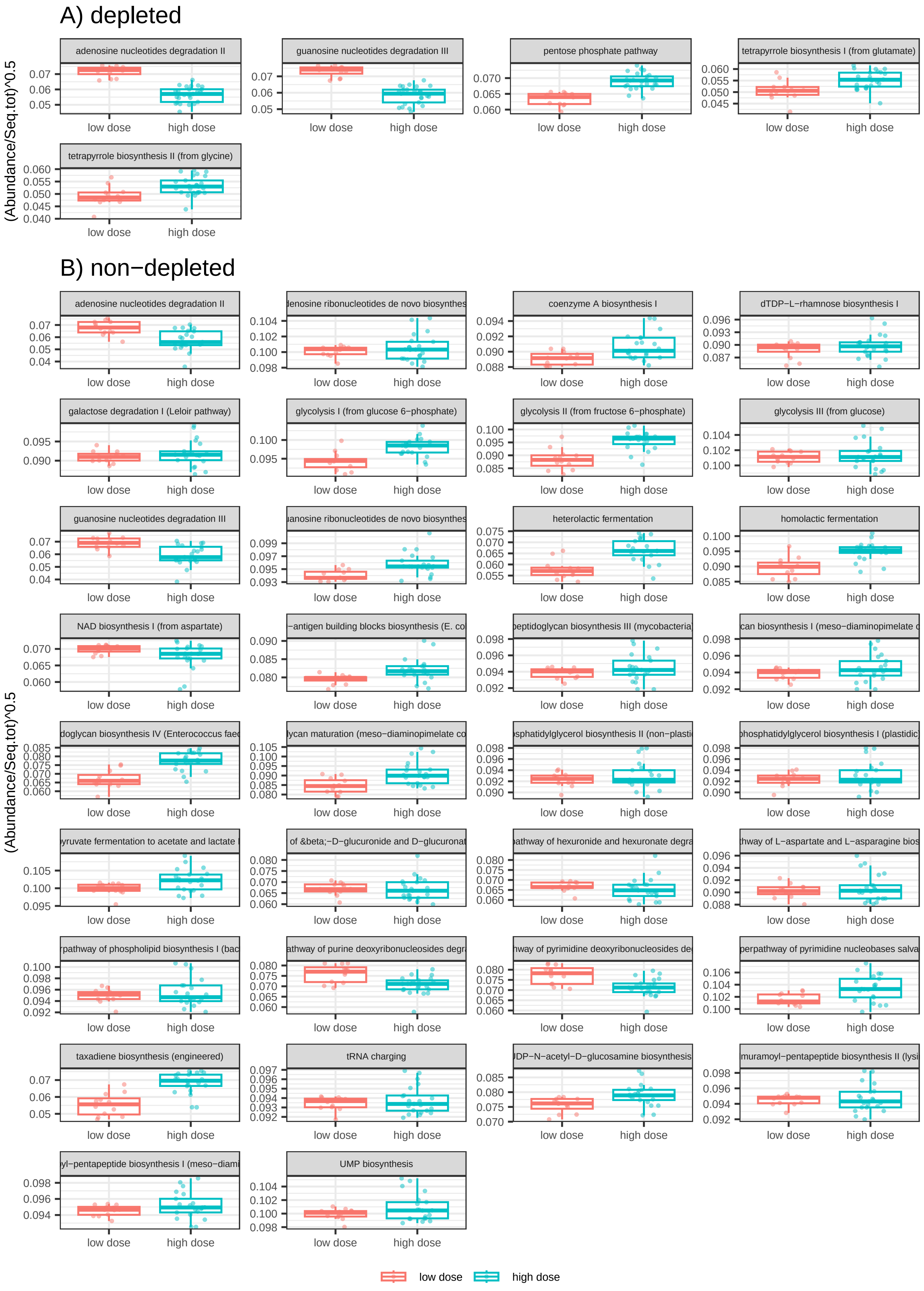

### Figure S7

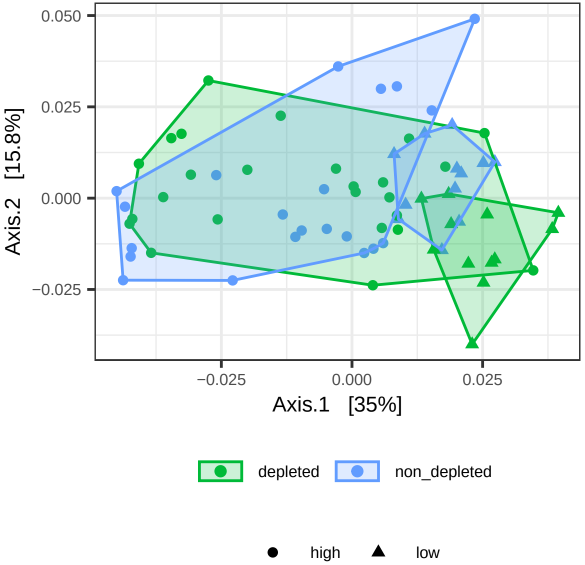

### Figure S8

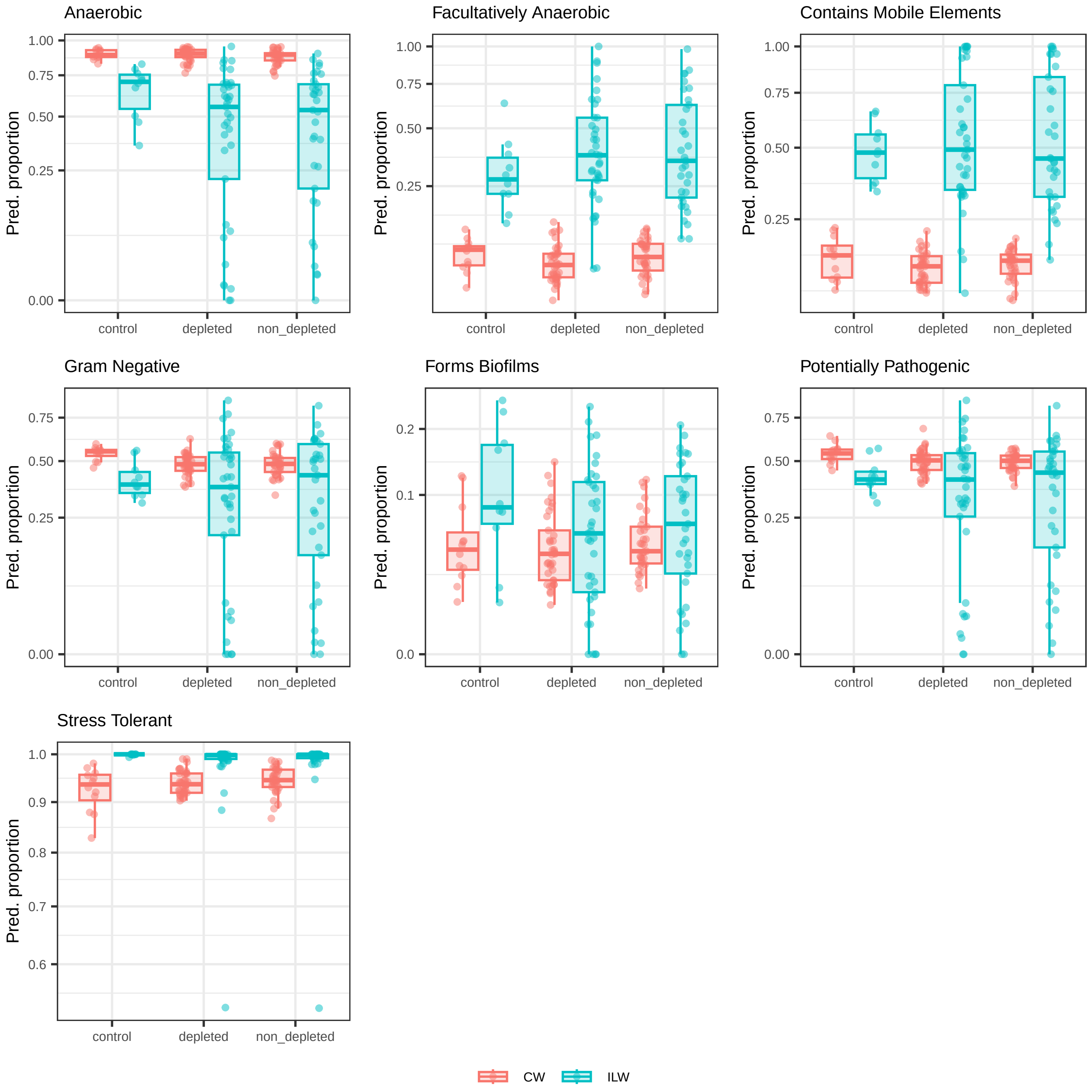
